# Recovering individual haplotypes and a contiguous genome assembly from pooled long read sequencing of the diamondback moth (Lepidoptera: Plutellidae)

**DOI:** 10.1101/867879

**Authors:** Samuel Whiteford, Arjen E. van’t Hof, Ritesh Krishna, Thea Marubbi, Stephanie Widdison, Ilik J. Saccheri, Marcus Guest, Neil I. Morrison, Alistair C. Darby

## Abstract

**Background:** Recent advances in genomics have addressed the challenge that divergent haplotypes pose to the reconstruction of haploid genomes. However for many organisms, the sequencing of either field-caught individuals or a pool of heterogeneous individuals is still the only practical option. Here we present methodological approaches to achieve three outcomes from pooled long read sequencing: the generation of a contiguous haploid reference sequence, the sequences of heterozygous haplotypes; and reconstructed genomic sequences of individuals related to the pooled material.

**Results:** PacBio long read sequencing, Dovetail Hi-C scaffolding and linkage map integration yielded a haploid chromosome-level assembly for the diamondback moth (*Plutella xylostella*), a global pest of *Brassica* crops, from a pool of related individuals. The final assembly consisted of 573 scaffolds, with a total assembly size of 343.6Mbp a scaffold N50 value of 11.3Mbp (limited by chromosome size) and a maximum scaffold size of 14.4Mbp. This assembly was then integrated with an existing RAD-seq linkage map, anchoring 95% of the assembled sequence to defined chromosomal positions.

**Conclusions:** We describe an approach to resolve divergent haplotype sequences and describe multiple validation approaches. We also reconstruct individual genomes from pooled long-reads, by applying a recently developed k-mer binning method.

## Introduction

Technical and analytical advances in genomics have dramatically improved the achievable standard of genome projects. The amount of high molecular weight (HMW) DNA required to perform long-read sequencing has reduced significantly and has been accompanied by a steady increase in sequencing read lengths and read accuracy(1). Nevertheless, the reconstruction of these data into accurate haploid (or phased diploid) genome representations still poses a challenge. In particular, current algorithms are not fully robust to differences in heterozygosity between loci and across clades (2). Importantly, the differences in type and structure of heterozygosity are the consequences of unique evolutionary histories. Therefore, algorithms that cannot solve heterozygous regions of varying divergence in a generalised way across the tree of life may lead to systematic biases in our understanding of genomic variation within and between species.

A particularly challenging aspect of genome assembly is the detection of highly divergent regions (HDRs) (2), which often cannot be determined as allelic within the assembly process and requires supervised analysis (3). HDRs are likely to be biologically important and may indicate regions experiencing diversifying selection, for example the MHC locus in humans (4). Failure to properly resolve them could impact downstream analyses, particularly the detection of balancing selection and overdominant loci. For most heterozygous loci, reads are long enough to span haplotypes, haplotype divergence is low and repetitive sequence content is low, causing assembly graphs to form simple “bubble” structures (5,6). These simple occurrences can be partitioned and subsequently collapsed or phased (2,5,7). Alternatively, pedigree information, such as trios, can be leveraged by either partitioning F1 read data *a priori* (8,9) or integrating the pedigree information into the assembly graph itself (10). These latter approaches are robust to HDRs, and in organisms with relatively low repeat content, the reconstruction of fully phased diploid chromosomes is eminently achievable.

Numerous initiatives have begun, or are proposing to generate genomic data for huge numbers of organisms across the tree of life. Pedigree sequencing is too reliant on obtaining or rearing mating pairs to be a feasible approach for these projects. Similarly, for the multitude of organisms with small amounts of extractable HMW DNA per individual, it may not be possible to avoid pooled sequencing. The likely best-case scenario for many arthropod samples is a small number of individuals from the same population. Here we analyse a pooled long-read dataset for the diamondback moth *P. xylostella* and show that additional contiguity information can be retrieved from such data, post-genome assembly. Furthermore we test the ability of k-mer binning methods to partition haplotypes from this pool using closely related but non-parental data (fig. 1). These methods were then critically assessed using sequencing coverage, gene content, shared-synteny and comparative k-mer coverage.

**Figure 1:**
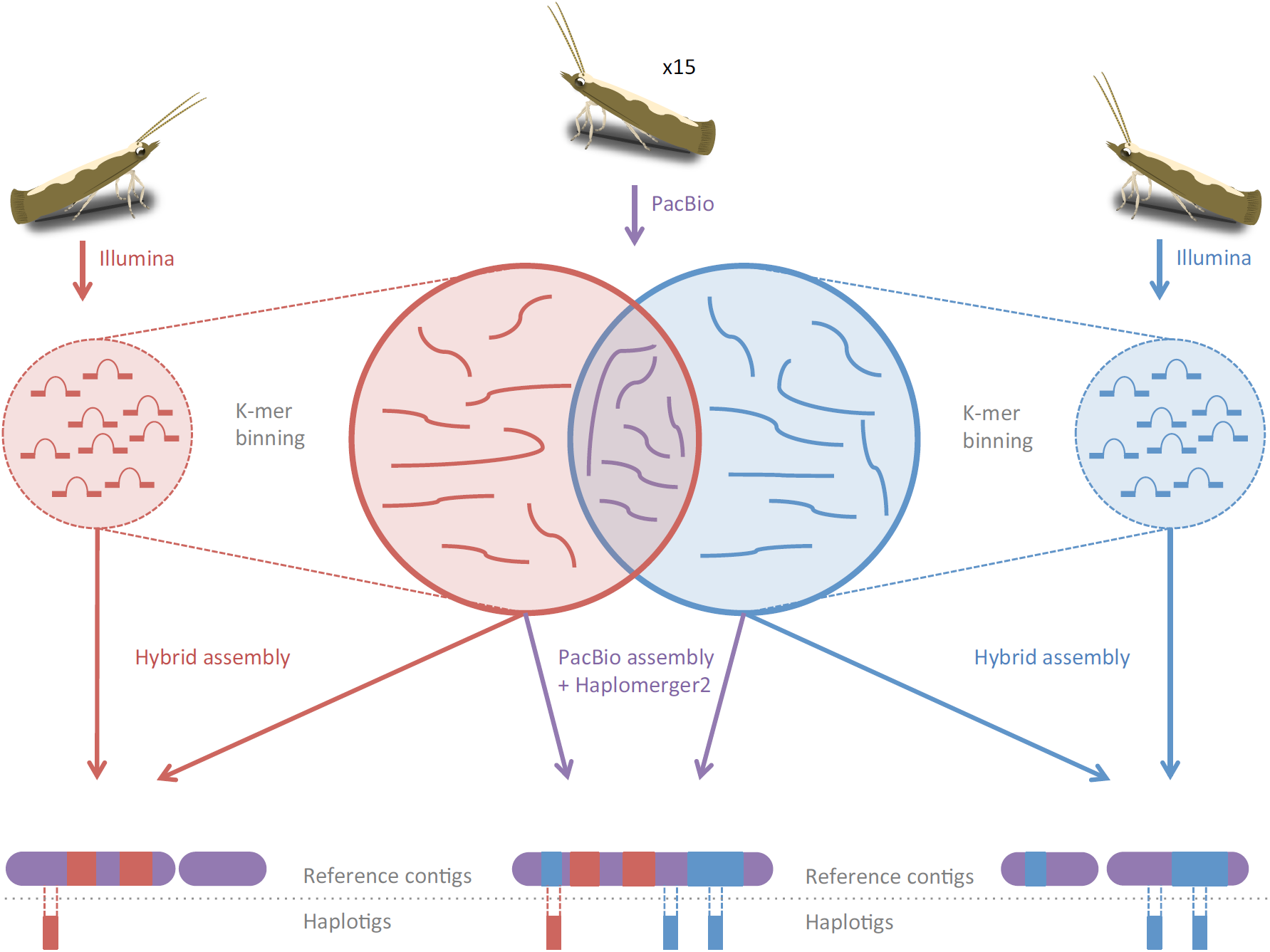
Strategies for assembling pooled long-read data. The coloured circles surrounded by the dashed line represent paired-end sequencing reads. The connection of these circles to the larger solid coloured circles represents the ability to use the k-mers of paired-end reads to identify long-reads of the same haplotype from pooled long-read sequencing of related individuals. These haplotype-assigned long reads can then be assembled alongside the short-reads using hybrid assembly methods, to approximately reconstruct an individual. Alternatively the pooled long-reads can be assembled using haplotype-conserving parameters, these haplotypes can then be resolved using a post-hoc alignment method. The ideograms at the bottom of the graphic show how the different methods capture heterozygosity differently in the resulting assemblies.

*Plutella xylostella* was the subject of a major genome sequencing effort, culminating in the publication of an assembly in 2013 (GCA_000330985.1)(11). The assembly strategy utilised the sequencing of fosmids in order to mitigate the short read lengths of Illumina sequencing. The authors report extensive structural variation based on alignments between their assembly and both the fosmids and a previously sequenced BAC (GenBank accession GU058050). The genome of *P. xylostella* therefore represents two distinct challenges to current long-read assembly methods, namely a large proportion of structural variation and a small amount of extractable DNA per individual. Our study includes the additional challenge of sequencing the heterogametic sex.

## Methods

### Sequencing material

Starting material was provided by Oxitec Ltd. (Abingdon, U.K.) from a lab colony that has been continuously cultured on artificial diet(12). For short-read sequencing DNA was extracted from a single male and single female individual. For the long-read sequencing, several lines were inbred in parallel by mating sib-sib pairs each generation. Many lines were eliminated due to severe inbreeding depression and one line that lasted for 7 generations was selected for genome sequencing. For long-read sequencing, DNA was extracted from 15 sisters of the final partially inbred generation.

### Library construction and sequencing

The pooled DNA was sheared to 7Kbp or 10Kbp. A subset was size selected at 15Kbp on the BluePippin (Sage Science, Inc.). In total 66 SMRT cells were sequenced with P5-C3 chemistry on the RSII platform (Pacific Biosciences, Inc.). Reads were filtered according to subread length (>50), polymerase read quality (>75) and polymerase read length(>50). Extracted DNA from the individual male and female was sheared and used for individual libraries followed by 2x 100bp paired-end Illumina sequencing (Illumina, Inc).

### Genome assembly parameters

FALCON genome assembly was performed using the parameter set recommended for the most appropriate similar organism, *Drosophila melanogaster*. A number of Canu (v1.4 and v1.5) parameter combinations were tested to help reduce heterozygosity during assembly (supp. tab. 1). The final assembly version was generated according to the following procedure: Assembly was conducted with Canu (version 1.5) using parameters recommended for conserving haplotypes (corOutCoverage=200 correctedErrorRate=0.040 “batOptions=-dg 3 -db 3 -dr 1 -ca 500 -cp 50”). The resulting assembly was polished twice with Quiver followed by an additional polishing step using Pilon (version 1.21) with female Illumina libraries mapped with BWA MEM (0.7.5a)(13–15). Mis-assemblies were identified and located using syntenic blocks (computed by SyMap version 4.2) corresponding to different *Bombyx mori* chromosomes(16,17). Mis-assembled contigs were then broken manually at the identified locations, referring to aberrations in PacBio and Illumina read mapping using IGV (version 2.3.40)(18)(Thorvaldsdóttir et al., 2013). Screening for contamination was performed with BlobTools (version 0.9.19) to the genome and mapped Pacbio reads generated after the first round of Quiver polishing(19).

### Haplotype merging

The polished genome was masked using a custom database of Lepidoptera repeats extracted with the queryRepeatDatabase.pl script included with RepeatMasker (1.323) in combination with a *P. xylostella* repeat library downloaded from LepBase (archive version 4)(20,21). A species-specific scoring matrix was inferred at 95% identity using the lastz_D_Wrapper.pl script included with Haplomerger2(22). The masked genome and scoring matrix were then used to run scripts B1-B5 of the Haplomerger2 pipeline (version 20161205)(22). Possible mis-joins were identified using the same synteny-based method described previously. Mis-joined contigs and W-chromosome candidates were identified in the Haplomerger2 output files and prevented from merging.

### Scaffolding

The preparation and analysis of HiC and Chicago libraries were performed by Dovetail LLC. using pools of starved larvae. In brief, Chicago libraries were performed as described by Putnam *et al.*, 2016. HiC libraries were prepared as described by Kalhor *et al.*, 2012. Both library preparations used the restriction enzyme DpnI for digestion after proximity ligation. Scaffolding was performed by running HiRise, first using the Chicago data followed by a second iteration using the HiC data (23).

### Chromosome assignment

The RAD-seq linkage map generated by Baxter *et al.* (24) was used to order and orientate sequences into putative chromosomes. The linkage map was generated *de novo* using STACKS (25). The consensus RAD-tag loci were mapped to the genome with Stampy (version 1.0.2)(26). Reads were filtered by mapping quality at two thresholds (1 & 30), forming the basis of high and low confidence assignments. RAD-tag associated contigs were also mapped using the same procedure and a mapping quality cutoff of 70. The mapped marker and sequence data was integrated using Chromonomer (version 1.03)(27).

### Haplotype read and haplotype assembly mapping

Reads were binned into haplotypes using the TrioCanu program included with Canu (version 1.7). Binned reads were mapped to the finished genome with BLASR and filtered with samtools by mapping quality (>30). Binned reads were assembled together with the paired-end reads using MaSurCA (version 3.2.6b) using recommended parameters, without linking mates (28). The original assembly and hybrid assemblies were mapped back to the finished reference with Mummer (v 3.23)(29).

### Validation procedures

Canu-corrected long reads, the female paired-end short reads and the polished Canu assemblies before and after Haplomerger2 were all decomposed into 21-mer hashes using jellyfish (version 2.2.0). The comparison tables were then generated using the comp program included in the k-mer analysis toolkit (version 2.3.1). BUSCO results were generated with BUSCO (version 3) using the parameters “-m genome --long --limit 4 -sp heliconius_melpomene1” in combination with the Endopterygota odb9 database. Synteny was examined with SyMap (version 4.2).

## Results

### Assembly

After filtering 2,655,788 PacBio subreads remained (mean subread length=7,301bp, N50=10,398bp, total bases 19.4Gbp). Almost all assembly strategies resulted in an over-inflated genome size (fig. 2), which was suspected to be due to redundant alleles assembling as independent contigs (termed haplotigs)(5). Both Canu and FALCON assemblers performed similarly (fig. 2).

**Figure 2:**
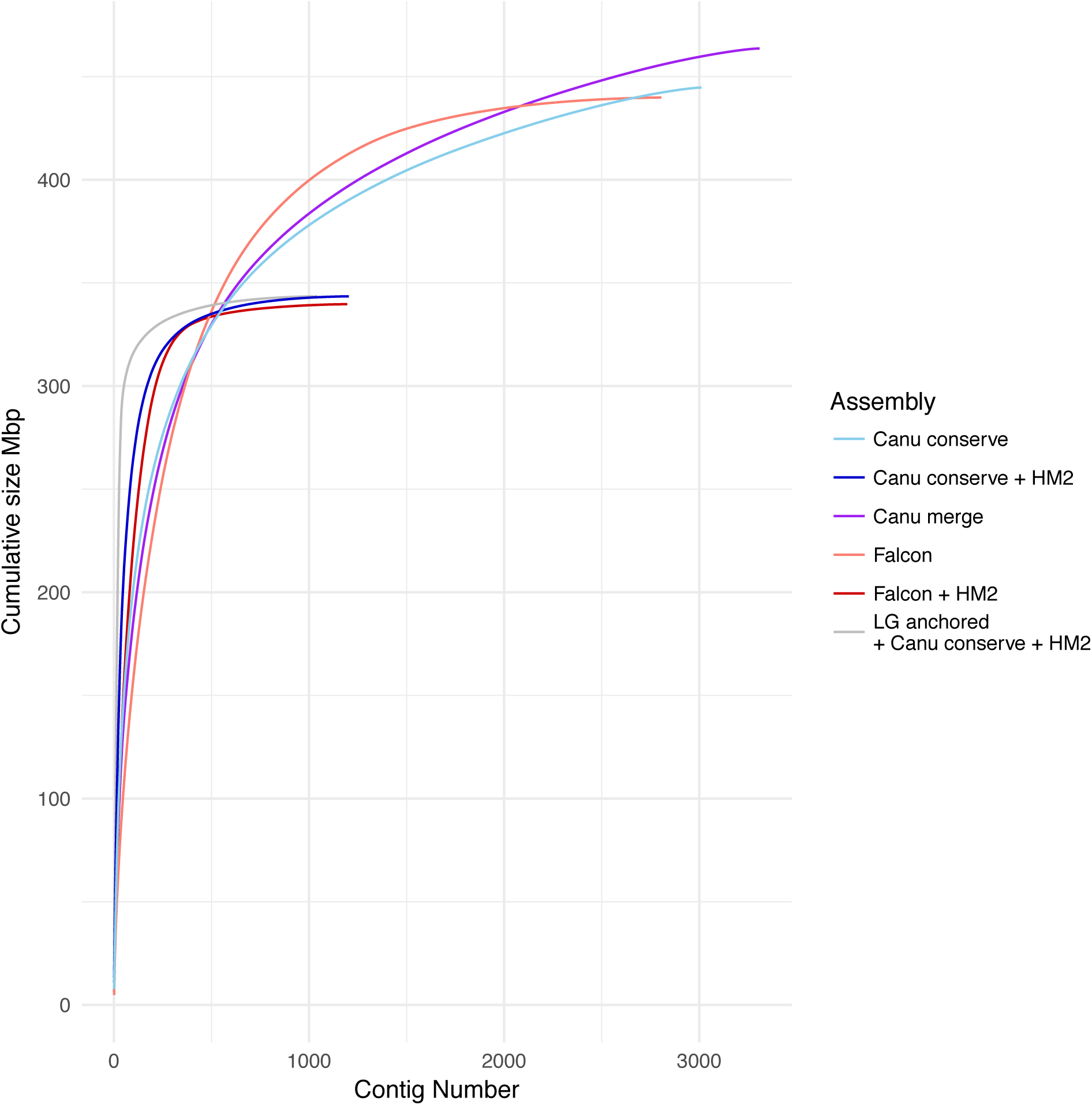
Cumulative size plots of Canu and Falcon assemblies before and after haplotype merging. The application of post-hoc haplotype merging results in a number of positive outcomes, in particular the adjustment of the total assembly size much closer to the genome size estimated by flow-cytometry(24). In addition, the total number of contigs is dramatically reduced and the increased curvature suggests that some contigs are tiled into longer contiguous sequence.

A range of parameter alterations for Canu (v1.4) were made in an attempt to improve assembly contiguity but ultimately had little influence (supp. tab. 1). Increasing the number of corrected reads available for construction of the assembly graph is theoretically useful as coverage is reduced for highly heterozygous regions; this is exacerbated for the Z-chromosome, which already has half the autosomal coverage. Increasing the coverage of corrected reads improved contiguity relative to default parameters (NG50 +31.7Kbp); however max contig length decreased (−479.8Kbp). Increasing the error-rate parameter theoretically allows divergent alleles to be merged during assembly. When the error-rate parameter is increased, reads from regions containing heterozygous SNPs can still be overlapped (in effect SNPs can be treated as errors). This does not however address more troublesome structural heterozygosity and it is likely to introduce spurious edges in the assembly graph(15). Increasing the error-rate reduced contiguity relative to default parameters (NG50 -103.5Kbp), max length increased (+3.2Mbp). Increasing a third parameter “-utgGraphdeviation” is similar to the previous parameter in that it enables alleles containing divergent SNPs through more permissive alignments and bubble collapsing during assembly graph construction. Setting utgGraphDeviation at 10 or 28, resulted in virtually identical assemblies to that obtained with default parameters. Two additional parameter sets were tested for a later version of Canu (v1.5) following recommendations for conserving haplotypes (parameter set 7, supp. tab. 1) and merging them (parameter set 8, supp. tab. 1) during assembly. The haplotype merging parameters appeared to negatively impact the assembly of the longest sequences and counter intuitively resulted in a larger total size and number of contigs.

The suggested FALCON parameters for *Drosophila melanogaster* were used for *P. xylostella.* Aside from being the most closely related organism for which parameters are provided, they have reasonably similar sized genomes, though they differ in chromosome structure (lepidopterans are holocentric). Despite FALCON being designed explicitly for resolving heterozygosity, the initial assemblies were relatively similar to Canu (v1.5) assemblies in that the resulting total sequence remained much larger than the expected genome size. Successful heterozygosity resolution should result in both decreased total length and increased contiguity, without a large impact on the gene space. Increasing the pre-assembled (pread) read length, decreased the total genome size by 22.2Mbp and NG50 by 40.4Kbp. The decrease in contiguity indicates that the decrease in genome size is likely to be the result of some haplotypes no longer having sufficient coverage to assemble. Ultimately Falcon and Canu produced comparable assemblies in terms of contiguity, completeness and total size. Post-assembly haplotype merging was performed on both. Results indicated that Canu responded more favourably to this procedure than Falcon (fig. 2). Interestingly, we found that the Canu assembly with haplotype conserving parameters was able to more effectively form a tiling path with Haplomerger2 than other assemblies, thereby achieving increased contiguity (Sup. fig. 3). The haplotype merged Canu assembly consisted of 1204 contigs, with a total assembly size of 343.5Mbp a contig N50 value of 2.9Mbp and a maximum contig size of 10.9Mbp. The Canu assembly was subsequently scaffolded with Dovetail Hi-C and HiRise data yielding an assembly consisting of 573 scaffolds, with a total assembly size of 343.6Mbp a scaffold N50 value of 11.3Mbp and a maximum scaffold size of 14.4Mbp. This assembly was then integrated with the RAD-seq linkage map, assigning 95% of the assembly scaffolds to linkage groups.

### Gene-informed heterozygosity assessment

The existence of paralogues within a genome can produce a similar signature to redundant alleles, making it difficult to distinguish the two cases without knowledge of the evolutionary history of the considered genes (31). The benchmarking universal single-copy orthologues (BUSCO) tool was developed to broadly assess the “completeness” of a gene-set and includes a measure of gene duplications (32). Importantly, the definition of BUSCO genes is based on an inference of evolutionary history (single-copy orthologues in >90% of sampled species) implying that duplications identified are either erroneous or rare (33). The result of BUSCO analysis of the haplotype-conserving *P. xylostella* assembly and the previous reference genome included a large number of duplicated genes. This is particularly striking when viewed alongside other lepidopteran genomes (Fig. 4). Our haplotype-conserving genome assemblies appeared to contain even higher levels of duplication than the previous reference, though it was unclear how much of this was the result of allelic redundancy. Examining how the identities of these duplicated BUSCOs intersect between the previous reference and the haplotype-conserving genome assembly, reveals very little overlap (Fig. 6). This could suggest that a large number of gene duplications in both datasets are in fact haplotype-induced artefacts, or population-specific duplications. Removal of redundant haplotigs from our initial assembly produces a BUSCO profile that appears typical for a lepidopteran genome (Fig. 4). The deduplication of BUSCOs in our genome assembly was examined using short read sequencing coverage. In the event that genes are erroneously deduplicated it is expected that the coverage of the gene would be approximately twice that of single copy genes. Our data does not show this pattern (Fig. 5), supporting the conclusion that BUSCO duplications in the haplotype conserving assembly are assembly artefacts.

**Figure 3:**
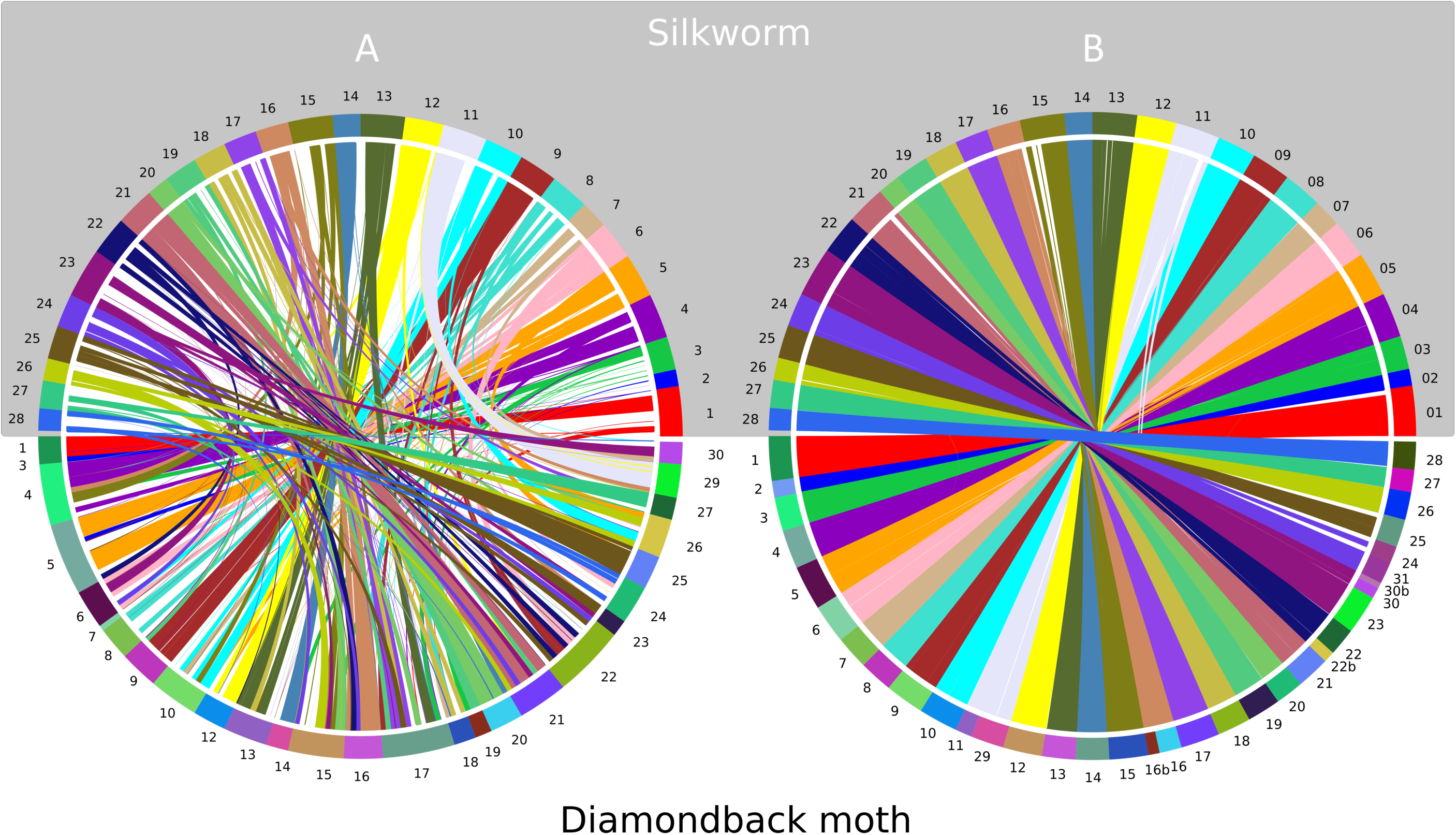
Shared-synteny between two *P. xylostella* genome versions and *B. mori*. The coloured bars around the top hemisphere of both plots represent the 28 chromosomes of *B. mori*. The coloured bars around the lower hemispheres of the plots represent the linkage groups produced by RAD-seq anchoring of (A) the previous *P. xylostella* reference genome and (B) the haplotype merged Canu assembly(11,24,30). Ribbons between the chromosomes indicate regions of shared-synteny identified by SyMAP.

**Figure 4:**
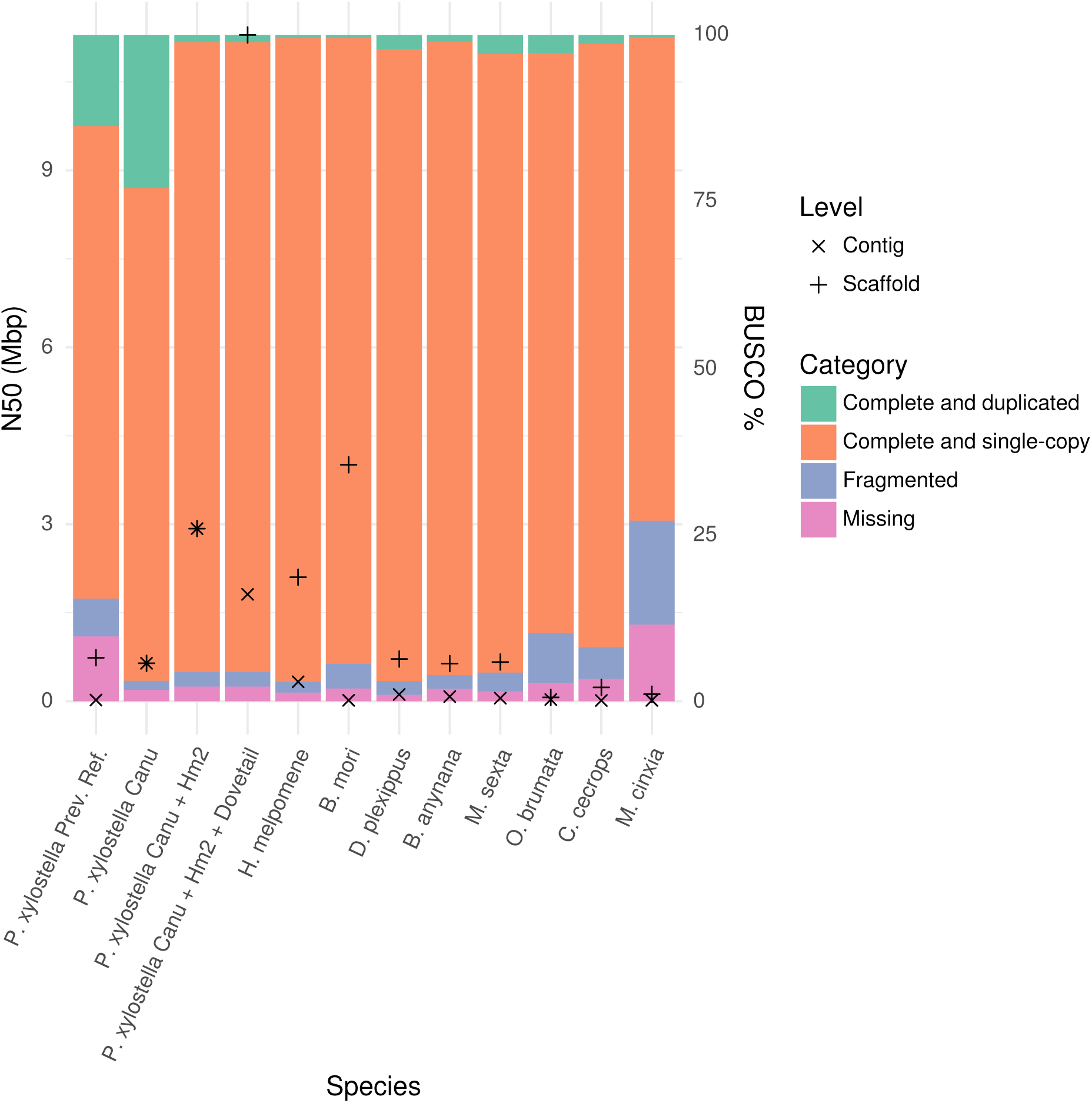
Quality comparison with other lepidopteran genomes. BUSCO profile and N50 at the scaffold and contig level are overlaid. The high levels of duplication in the previous *P. xylostella* reference and the haplotype conserving assembly is apparent, as is the effect of merging haplotypes which produces a profile similar to previously sequenced Lepidoptera (11,30,34–40).

**Figure 5:**
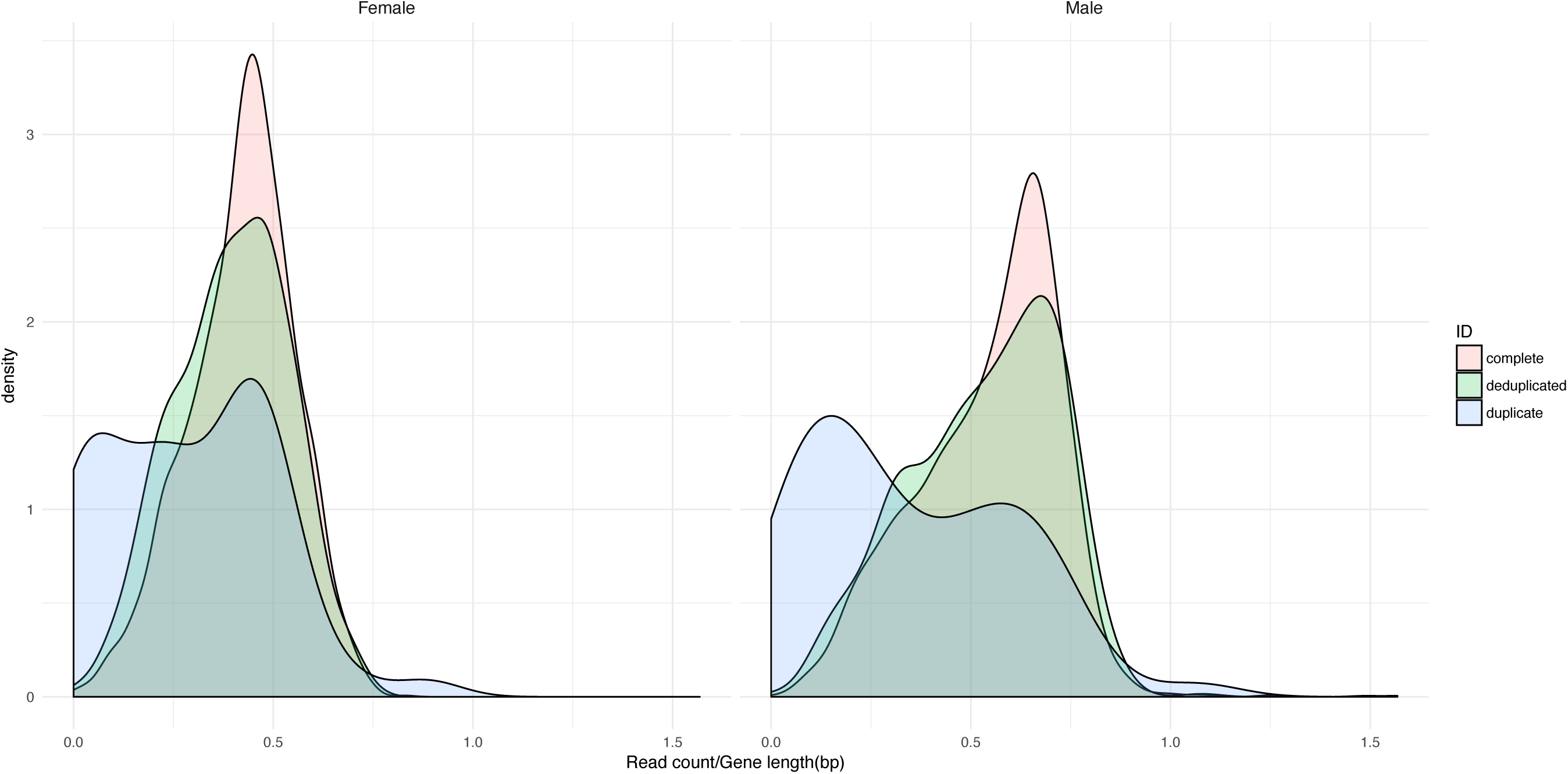
Coverage of BUSCOs in merged assembly. BUSCO coverage for mapped Illumina libraries. BUSCOs in the merged assembly are complete (single copy), duplicated or deduplicated (duplicated in the original assembly single copy after Haplomerger2). The erroneous collapse of real duplications would theoretically lead to a bimodal distribution for the deduplicated (green) genes, with one peak at approximately the same position as single copy genes (red) and an additional peak with two-fold higher coverage. The lack of this higher coverage peak supports the suggestion that redundant haplotypes are the basis for collapsed and merged sequence rather than segmental duplication.

**Figure 6:**
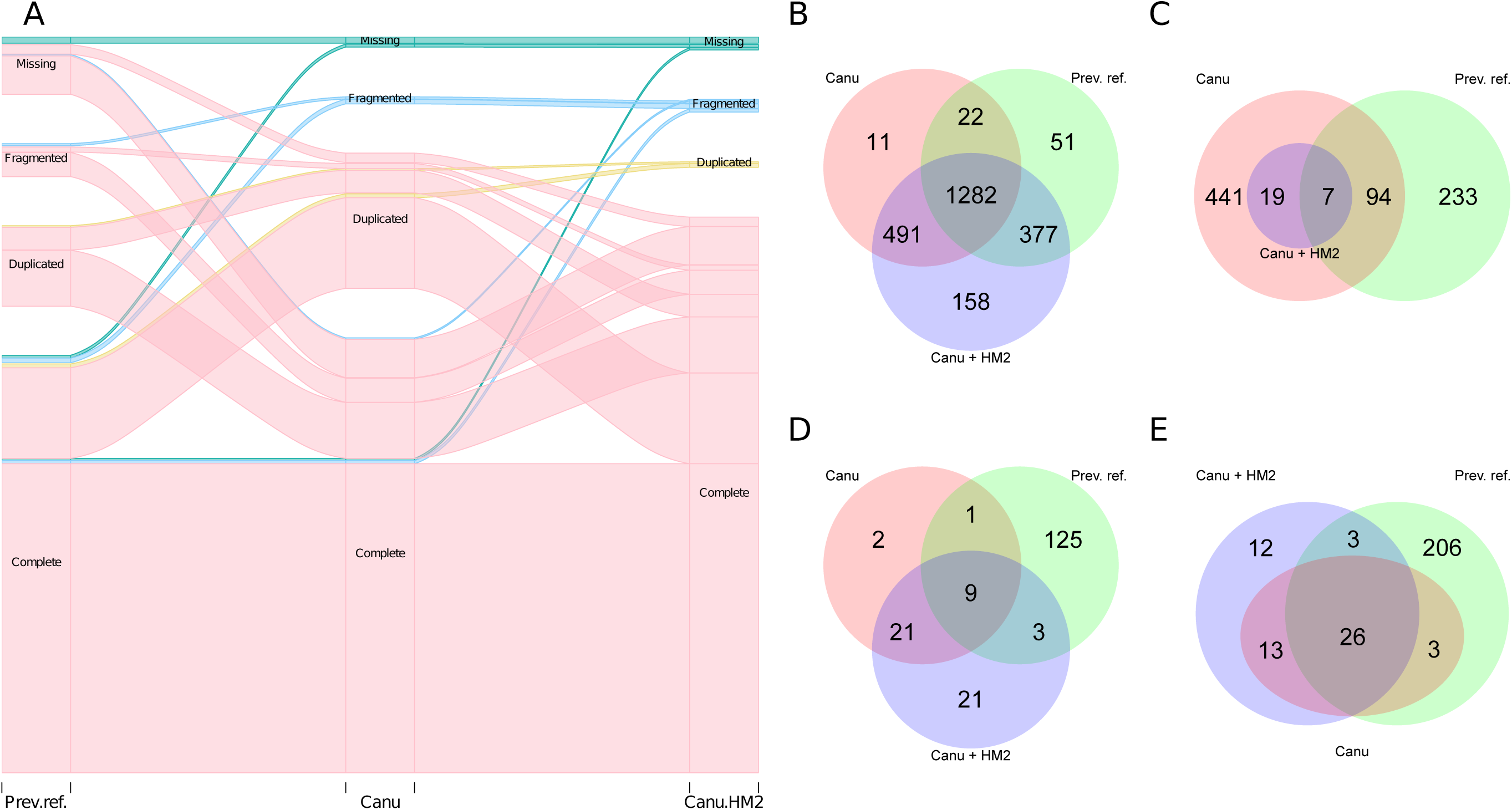
Between assembly BUSCO profiles. The alluvial plot (A) displays the movement of specific BUSCO genes between categories for the original and haplotype-merged Canu haplotype-conserving assembly, and the previous reference genome(11). The Venn diagrams (B,C,D,E) display the intersections of BUSCOs from each of the four categories (complete, duplicated, fragmented, missing), for the same three genomes.

### k-mer informed heterozygosity assessment

Using BUSCO genes to assess heterozygosity can yield indications, however only a subset of the total gene space is represented and it is possible that some gene duplications are true. An alternative strategy employed in the k-mer analysis toolkit (KAT), compares k-mer histograms from raw-reads against a genome assembly (41). Applying these plots to the haplotype-conserving and haplotype-merged assemblies, highlights a reduction in >1x assembly k-mers (Fig. 7). Additionally, the number of 0x assembly k-mers under the heterozygous peak increases, suggesting the effective removal of redundant sequence. The number of 0x k-mers under the homozygous peak also increased, which can be explained by the fact that the Illumina library was generated from a single female, whereas the assembly was generated from a pool of females, therefore alleles that are homozygous in the Illumina library are not necessarily expected to be homozygous in the assembly. Long-reads are difficult to leverage for this purpose due to their high error rate, though corrected long-reads, can give some indication (Fig. 7).

**Figure 7:**
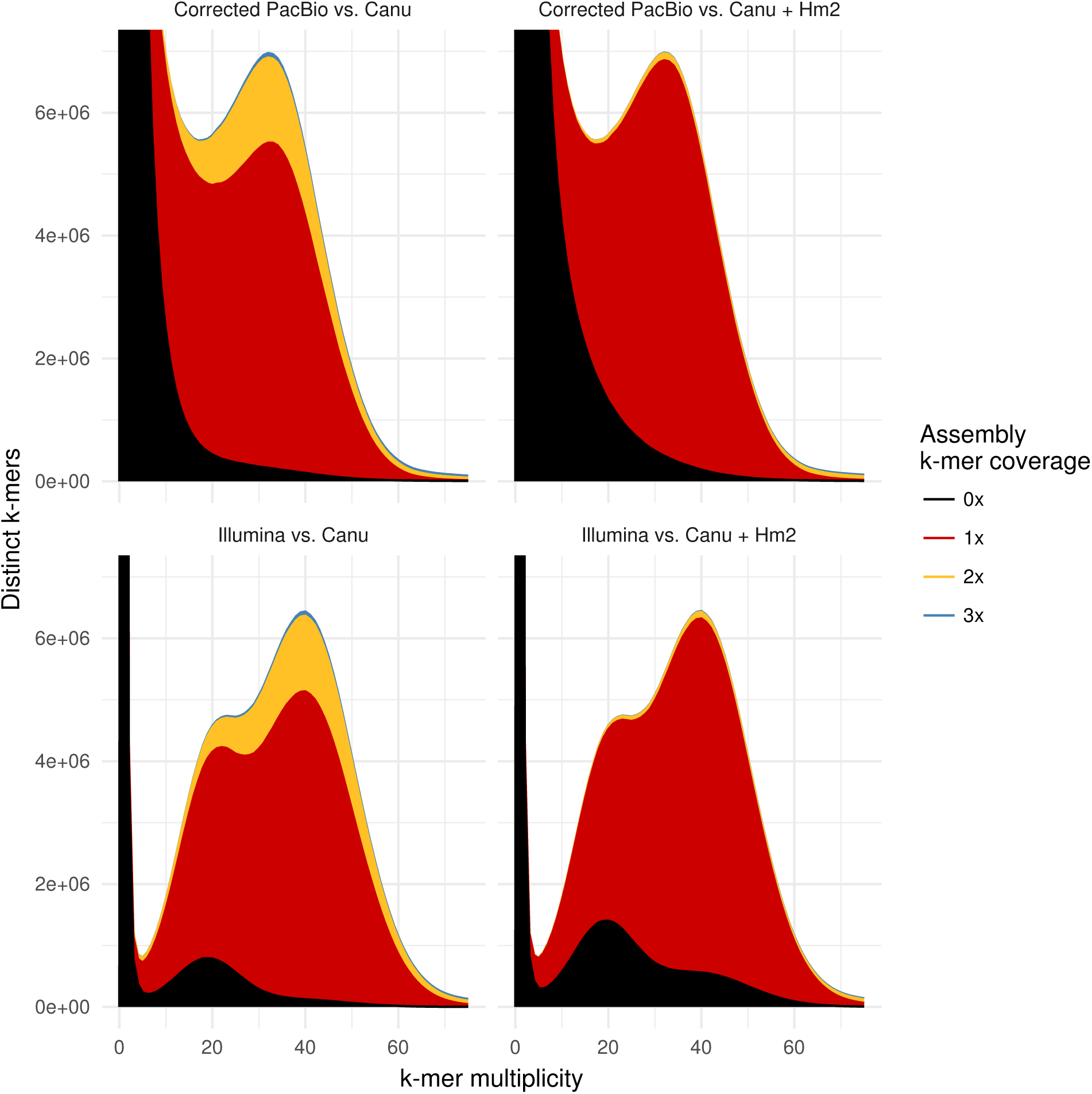
Stacked comparative k-mer histograms. Decomposition of corrected long reads produces the profile in the top panels, whereas decomposition of Illumina reads produces the profile in the bottom two panels. These profiles are then cross-referenced with k-mers occurring in the assembled data and the read profiles are “filled” according to assembly coverage. It should be noted that the PacBio reads were used to assemble the genome, whereas the Illumina reads were from a different single individual, therefore k-mers that are heterozygous in the assembly may be homozygous in the Illumina reads and vice-versa

### Heterozygosity mapping

Recent studies have shown that k-mer profiles can be used to determine the allelic origin of noisy long reads, this concept was originally applied in the context of an F1 hybrid, though it was recognised the concept could be expanded beyond a trio relationship (8). Utilising the same binning method with short-read libraries from an individual male and female, resulted in 1,625,290 female reads (12.9Gbp), 820,777 male reads (6.4Gbp) and 26,454 unknown (0.05Gbp) (fig. 8). Mapping these reads back to the haplotype-merged assembly, highlights switches between the conserved haplotypes (Fig. 9B). The k-mer classified PacBio reads were assembled *de novo* and with a hybrid approach (Zimin *et al.*, 2017). The latter strategy mitigates problems associated with the polishing requirements for long read assembly as the sequence is ultimately derived from short-reads, with long-reads acting as a guide. The hybrid assemblies appear to produce better results than *de novo* despite lower than recommended short read coverage (sup. fig. 2). The underlying assembly and the two hybrid assemblies were realigned to the haplotype merged and linkage group anchored assembly (fig. 9C). Overall the binned read assemblies only represent regions of the genome shared between the individual and sequences already present in the pool. The aligned hybrid female assembly covered 274.9Mbp (80%) of the genome and the male hybrid assembly 152.8Mbp (49%), though it should be noted that this excludes contigs that did not meet the similarity threshold (75%) or minimum contig length (5Kbp). Of these aligned portions the female hybrid assembly contained 334,059 SNPs and the male hybrid assembly 125,394. Interestingly, 18% of the original long read *de novo* assembly could not be realigned to the merged assembly and the number of SNPs was very low (140), whilst breakpoints were the largest feature category (alignment end not corresponding to sequence ends). Together these observations suggest that sensitive repeat-masking alignment chaining approach implemented by haplomerger2 is necessary to retrieve allelic relationships.

**Figure 8:**
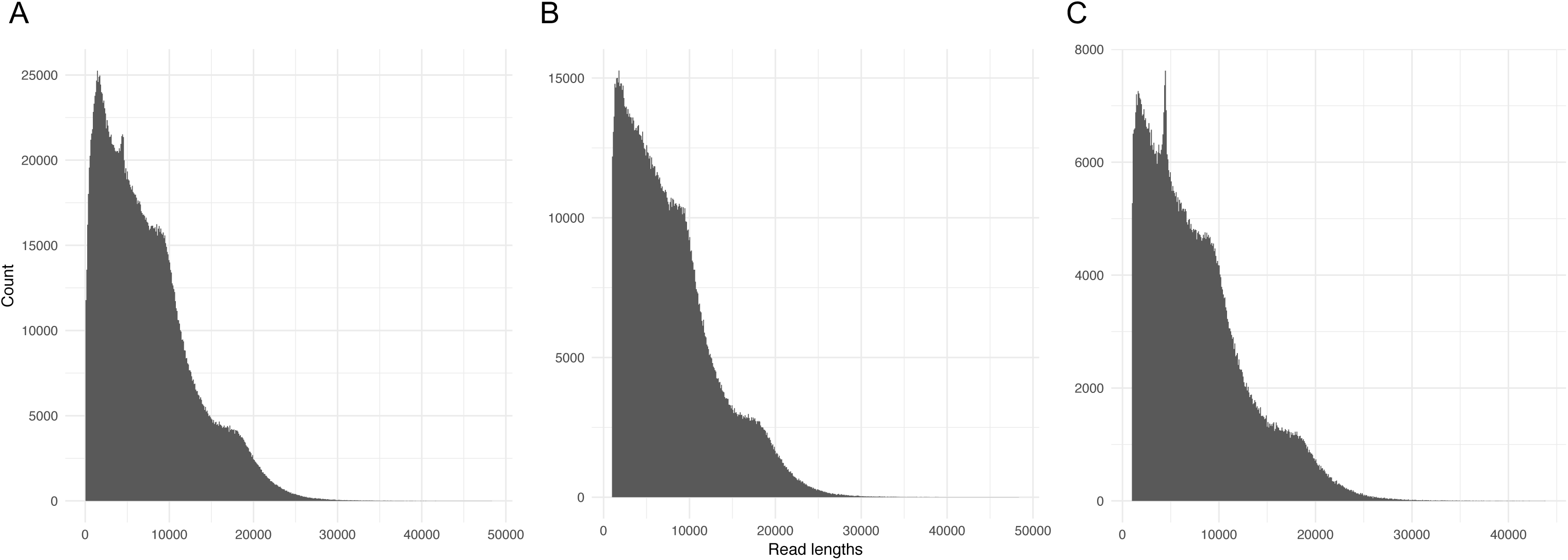
Histograms of filtered subread lengths. Total filtered subreads (A) and subreads after k-mer binning for female reads (B) and male reads (C).

**Figure 9:**
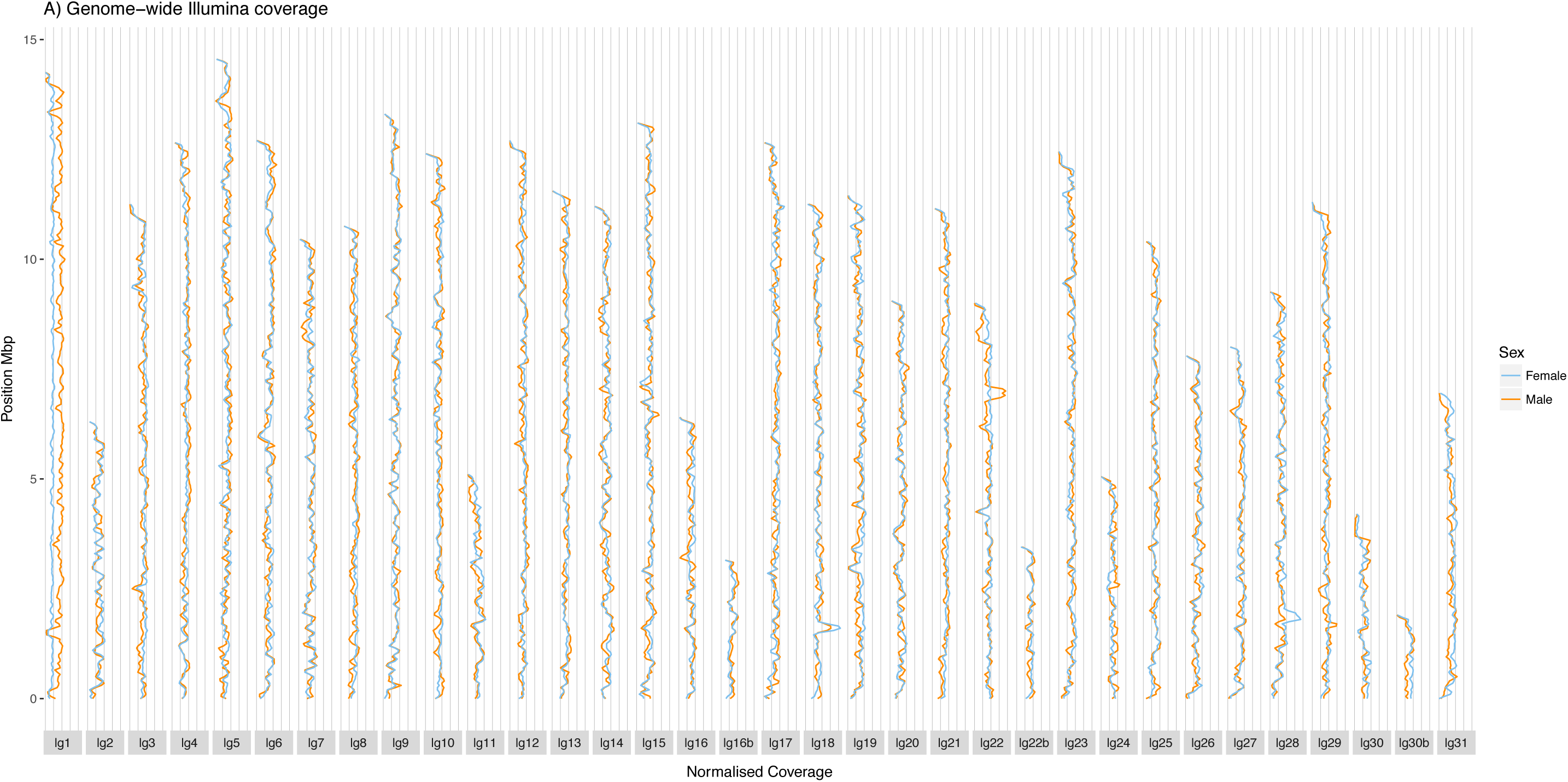
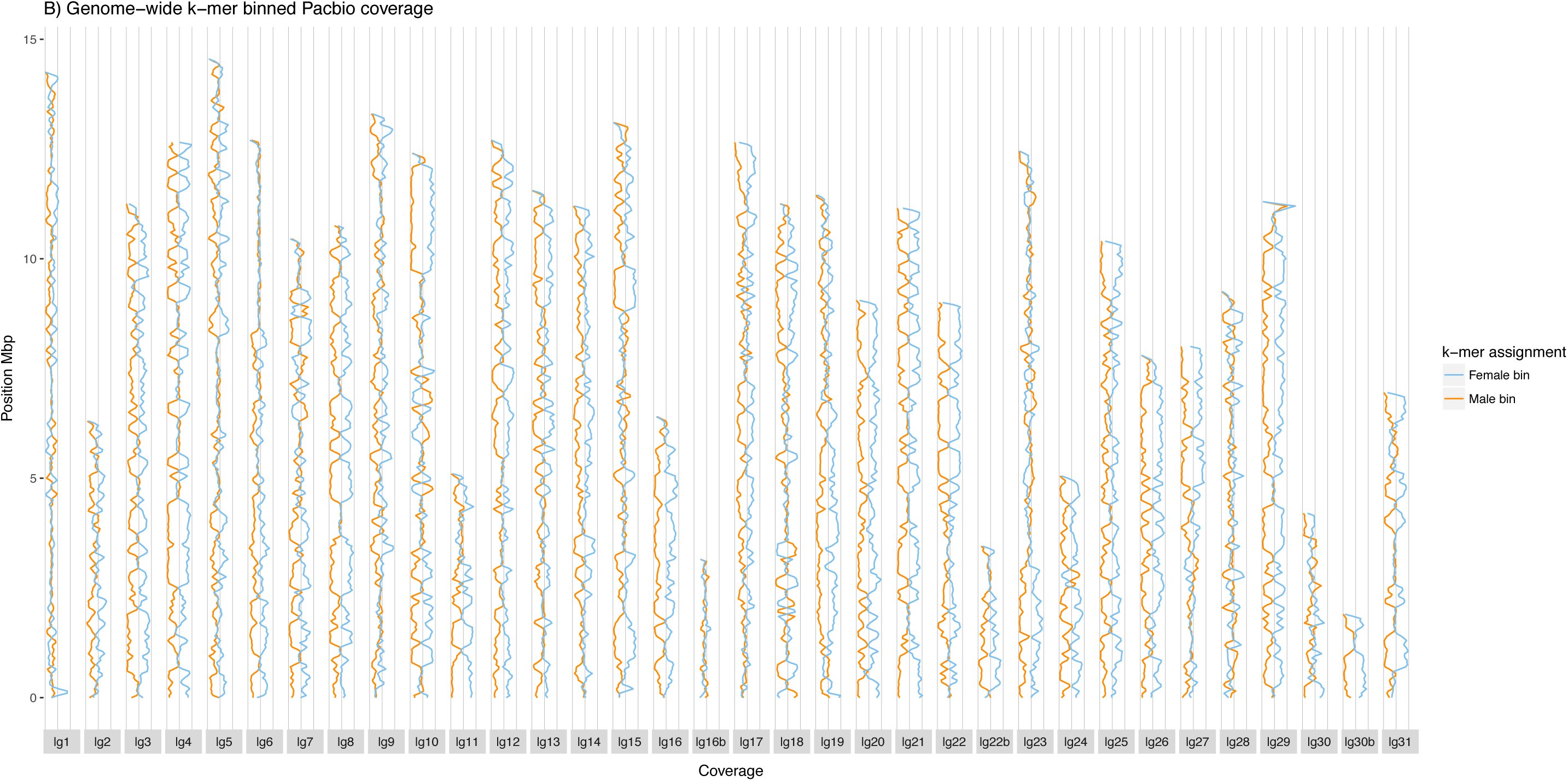
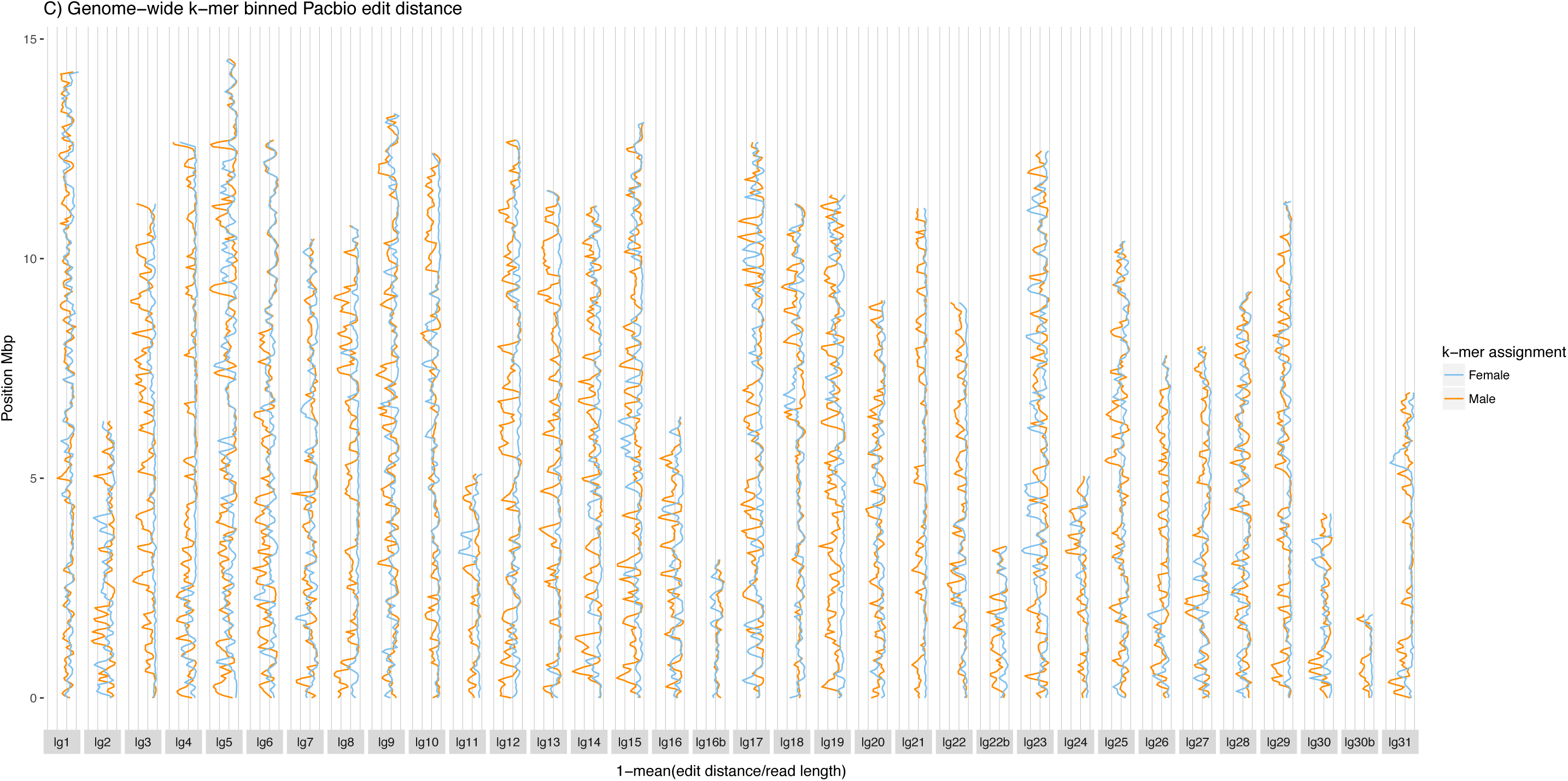
Assembly and read coverage plots anchored by linkage group positions. It is important to note that orientations are arbitrary for contigs with markers from a single mapping interval. (A) Illumina read counts normalised by total mapped counts in 100Kbp windows incremented in 50Kbp steps. The red line depicts the male library, whereas the blue line depicts the female library. X-axis lines denote intervals of 0.001. (B) PacBio read counts in 100Kbp windows incremented in 50Kbp steps. The red line depicts reads assigned to the male library k-mer bin, whereas the blue line depicts reads assigned to the female library k-mer bin. X-axis lines denote intervals of 250. (C) Depicts the edit distance calculated from the PacBio read alignments.

## Discussion

Gene family expansion was implicated by You *et al.*(11) in the ability of *P. xylostella* to rapidly evolve resistance to insecticides, as well as heterozygosity and transposon activity in the vicinity of detoxification genes. It is crucial to examine these conclusions in detail, as a propensity for gene duplication would distinguish *P. xylostella* from other lepidopterans (fig. 4). Our analysis of BUSCO profiles suggested that gene duplications reported in the previous reference genome could be haplotype-induced artefacts. Firstly, many genes duplicated in the previous reference are not duplicated in the long-read assembly and vice-versa (fig 6A & 6C). It is possible that this level of duplication may simply be the result of gene copy polymorphism between different populations, as described in *Heliconius sp.* populations (42); however the definition of BUSCO genes implies that such polymorphic duplication should be rare. When BUSCO genes are de-duplicated on the assumption of allelic redundancy, the coverage of these genes remains at a similar level to single copy genes (fig. 5).

Removal of allelic redundancy in the manner described here has been performed in other lepidopteran genome assemblies(34,35); however detailed validation of this process for *P. xylostella* was important, in the context of previous studies(11). Re-examination of the gene families described by You *et al.* (11) provides mixed support for their conclusions. In P450, GST and COE gene families, we find that the number of genes is much closer to *B. mori* than previously reported (supp. tab. 2). In contrast our data supports the finding that ABC genes are expanded in *P. xylostella* when compared to *B. mori* but still fewer than in the previous study. The possibility that these gene expansions are an innovation restricted to the East Asian populations or an artefact of different annotations cannot be discounted and requires more detailed study.

To meet the input requirements for PacBio library preparation it was necessary to pool multiple individuals. The expected heterozygosity was reduced without introducing W-chromosome variation, by extracting DNA from a pool of 15 sisters produced after 7 generations of sib-sib inbreeding. The amount of variation present in the source colony has not been fully quantified; however, this colony was itself derived from a different culture that was initially founded many years ago (12). It is therefore assumed that genetic should have already been relatively low in this population. Despite measures to reduce this variation further, it was still feasible that for any given autosomal loci there were potentially 4 unique alleles in the final inbred generation. The existence of HDRs within the final pool despite severe inbreeding of individuals from a colony derived from a long-term laboratory stock, could suggest the action of balancing selection or overdominant loci. Candidates identified by this study require further validation to partition the effects of drift from these aforementioned processes. The tiling path pattern seen in the original contigs suggests that in the absence of complete bridging reads, HDRs can be rescued (enhancing contiguity) using the sensitive alignment approach implemented in Haplomerger2 (22).

Comparing figures 9A and 9B further highlights the implications of heterozygosity for long-read assembly as opposed to short-read assembly. The mapping of Illumina reads shows only subtle deviations in coverage between the male and female libraries (fig 9A). For short-read data, haplotype divergence is expected to have a minimal impact on coverage as short reads only capture a small amount of variation and can often still be mapped in the presence of this variation. In contrast, long-reads contain a much greater amount of phased variation and therefore encode more haplotype specific information. This simple property is effectively utilised by the trio-binning method described by Koren *et al.* (8), providing an informed basis for haplotype phased assemblies. This study demonstrates that it may be possible to extend this concept further by extracting haplotype specific long-reads from a pool according to k-mers from short-read sequencing of an individual. In combination with the approach described by Zimin *et al.* (28), it is possible to use these “binned” long-reads in combination with the short-read data to assemble phased haplotypes, beyond what is achievable with short reads alone.

In parallel with this haplotype phasing approach, our realignment results suggest that general heterozygosity is not a particular challenge to current assembly methods. Instead it is the complexity of heterozygosity at a given locus that is the major challenge. Our results show that post-hoc methods can overcome this problem to a degree though it is not entirely error-free and requires validation. The haplotype merging method appeared to work particularly well with our data in terms of assembly contiguity and is amongst the best contig-level lepidopteran genome assemblies. In summation, the reconciliation of these two approaches yields both a highly contiguous genome assembly and a source of phased haplotype data. This study utilises a limited dataset, though we envisage that the conceptual approach is extendable to larger populations. For non-model heterozygous organisms for which extraction of HMW DNA is a limiting factor, this combination of pooled-long read sequencing and low coverage short-read sequencing may be an effective compromise.

The combination of pooled long-read sequencing with short read libraries is an effective and relatively cheap method of simultaneously generating genome assembly data and phasing variants from low-coverage short read data from single organisms within the same population. Our results indicate that whilst this strategy is sometimes pursued due to limitations on extracted DNA for long-read sequencing technology, the additional complexity introduced can be mitigated in a manner that preserves heterozygous sequence whilst maximising assembly contiguity.

## Data availability

Data submitted to the European Nucleotide Archive under project accession PRJEB34571 (see supp. tab. 3).

## Supplementary figures

**Supplementary figure 1:**
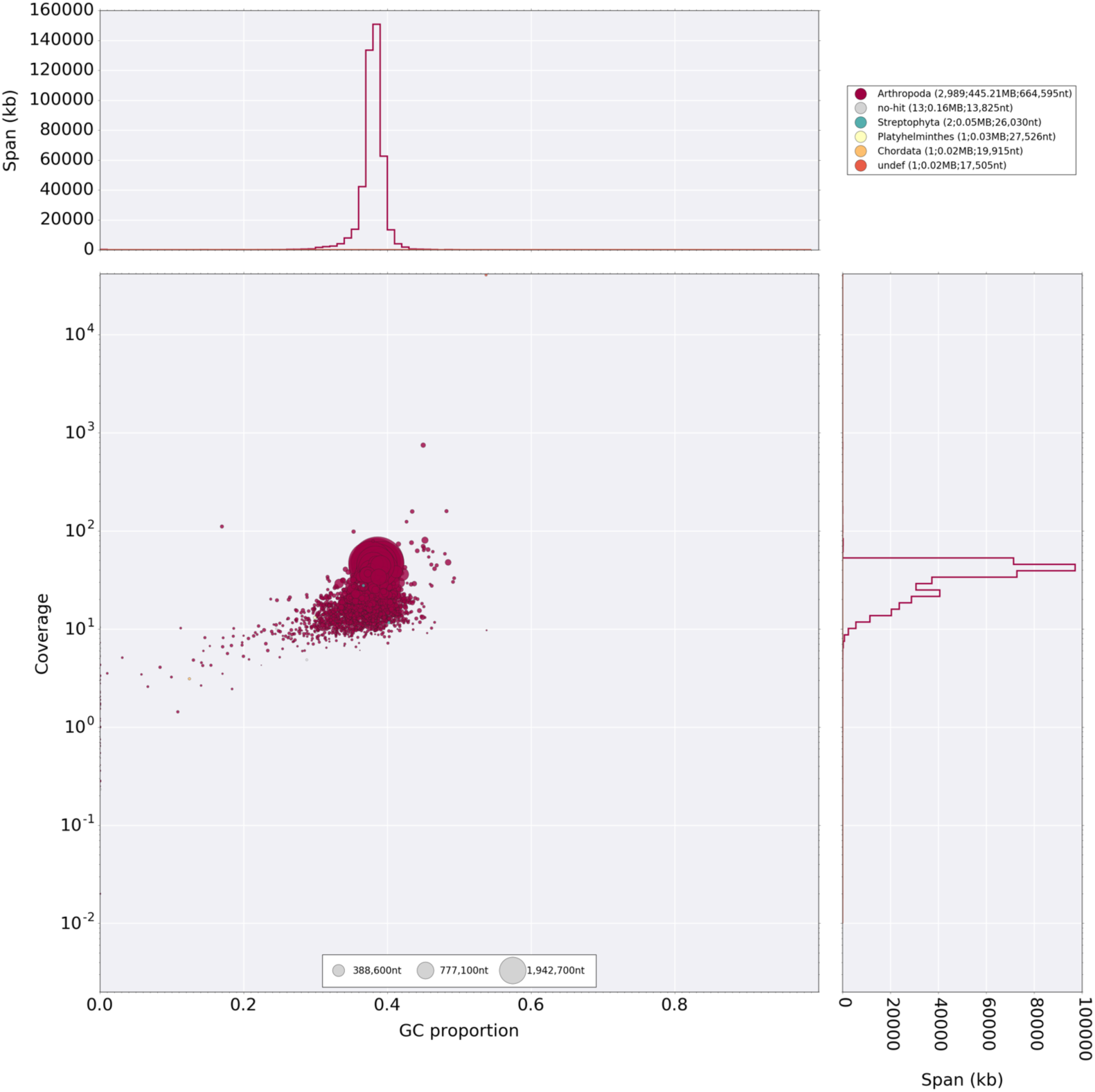
Blobtools contamination plot. PacBio reads mapped to the Canu haplotype conserving assembly from the second round of Quiver polishing were used to establish coverage. The plot shows no signs of contaminating DNA in the data.

**Supplementary figure 2:**
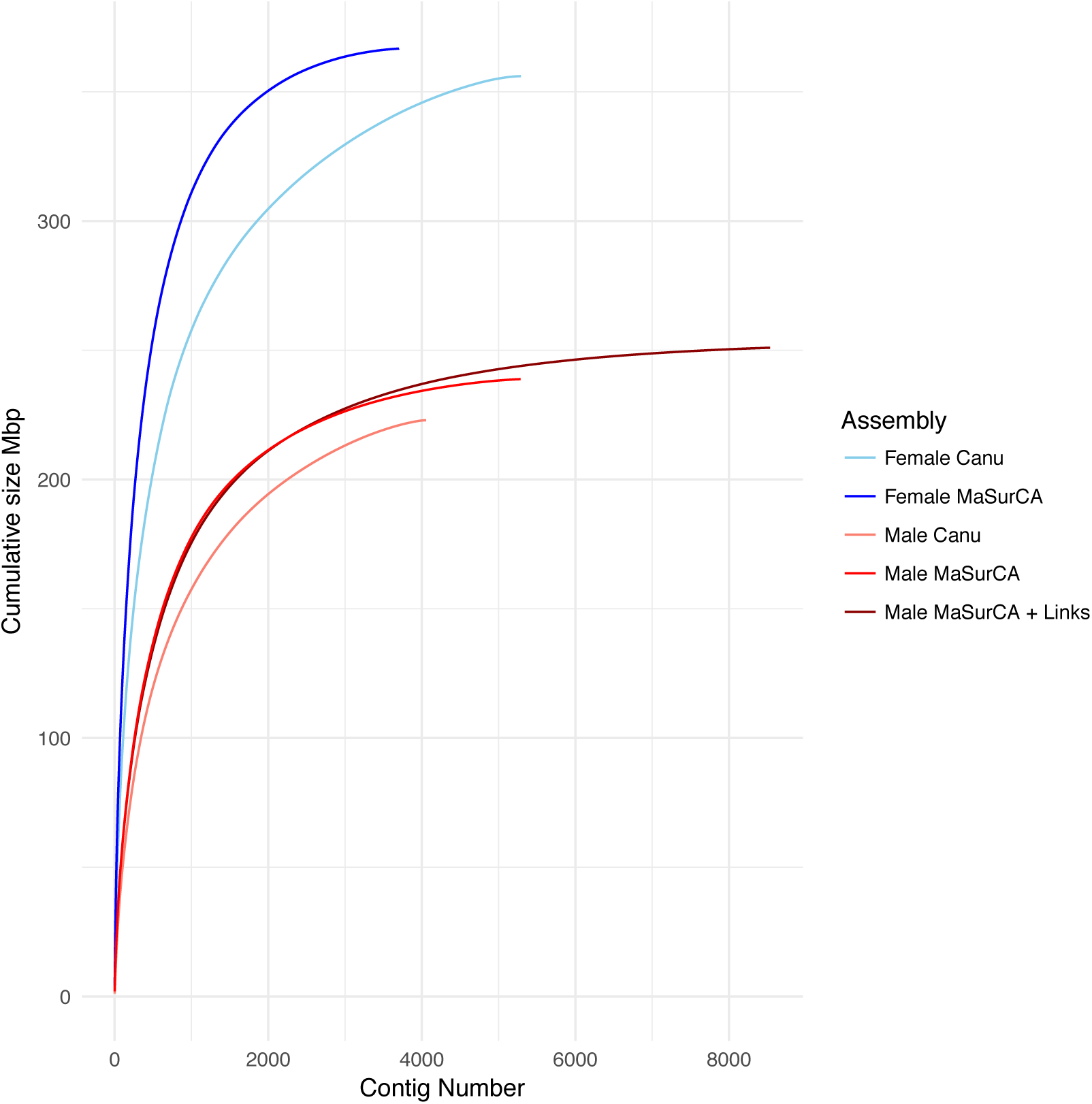
Cumulative size plots of genome assemblies of k-mer binned data.

**Supplementary figure 3.**
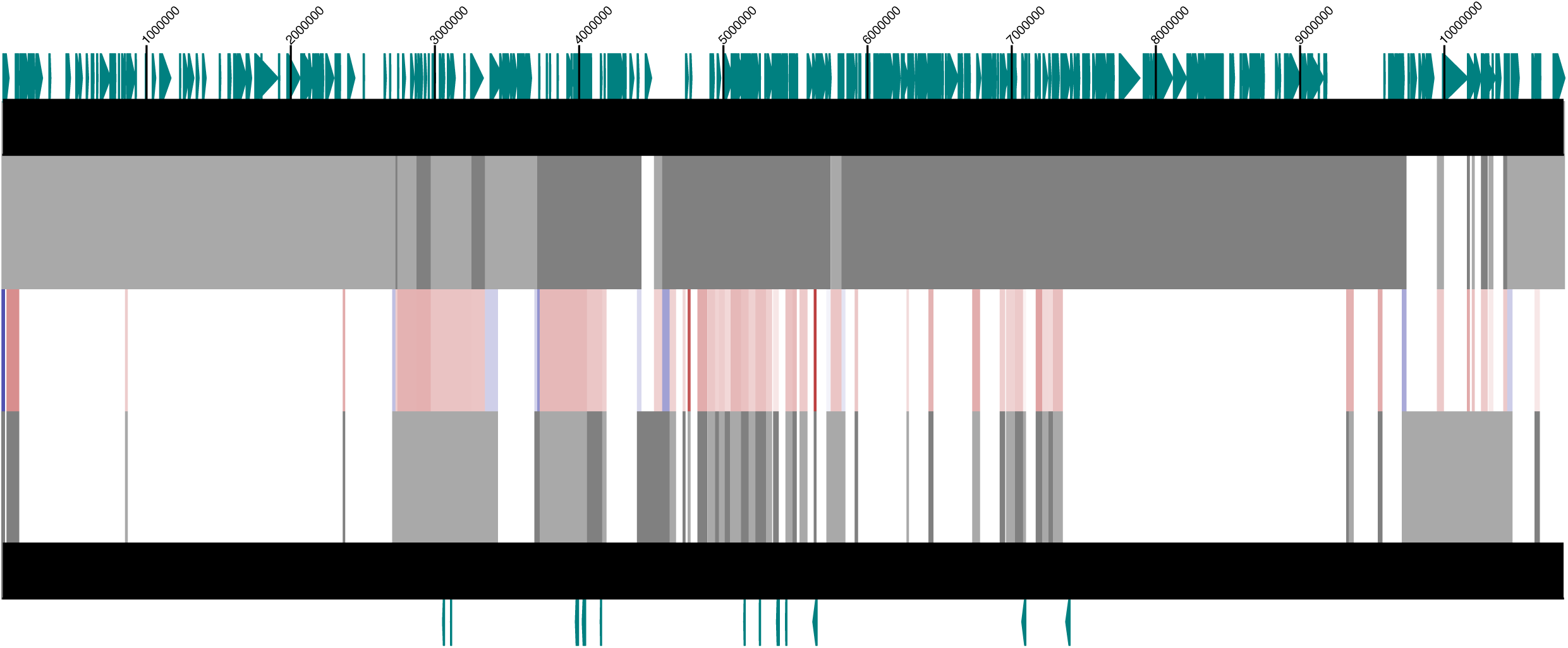
Tiling path of merged haplotypes: Grey boxes indicate the underlying polished Canu assembly (with conserve haplotype parameters). Red alignments indicate regions where contigs are collapsed within another larger contig by Haplomerger2, whereas blue alignments indicate regions where a contig forms a tiling path with another contig. The top track shows gene models along the merged contig, whilst the bottom track shows BUSCO genes that are de-duplicated by applying Haplomerger2

**Supplementary table 2:** Parameter details for Canu assemblies. This table provides a key of the various command line parameter alterations that were recommended by authors for resolving heterozygosity.

| Parameters | Canu version | Non-default parameters |
| --- | --- | --- |
| 1 | 1.4 | genomeSize=340m |
| 2 | 1.4 | genomeSize=340m -assemble<br>utgGraphDeviation=10 |
| 3 | 1.4 | genomeSize=340m -assemble<br>utgGraphDeviation=28 |
| 4 | 1.4 | genomeSize=340m errorRate=0.025 |
| 5 | 1.4 | genomeSize=340m corOutCoverage=100 |
| 6 | 1.4 | genomeSize=340m<br>corOutCoverage=100 errorRate=0.025 |
| 7 | 1.5 | genomeSize=340m corOutCoverage=200<br>correctedErrorRate=0.040 "batOptions=-dg 3<br>-db 3 -dr 1 -ca 500 -cp 50" |
| 8 | 1.5 | genomeSize=340m corOutCoverage=200<br>ovlErrorRate=0.15 obtErrorRate=0.15 |

**Supplementary table 2:** Re-estimation of previously reported gene family expansions. (N.R. = Not reported)

| Gene family | *P. xylostella* gene N.<br>(this study) | *B. mori* gene N.<br>(You *et al.*, 2013) | *P. xylostella* gene N.<br>(You *et al.*, 2013) |
| --- | --- | --- | --- |
| P450 | 86 (↓) | 84 | 93 |
| GST | 24 (↓) | 23 | 28 |
| COE | 71 (↑) | 76 | 66 |
| ABC | 61 (↓) | 51 | 94 |
| GSS | 2 | N.R. | 2 |
| Sulfatase | 8 (↓) | N.R. | 12 |

**Supplementary table 3:** Data accessions.

| Data | Accession |
| --- | --- |
| Conserved haplotype (diploid) assembly | ERZ1089753 |
| Haplomerger assembly | ERZ1089754 |
| Dovetail scaffolded assembly | ERZ1089755 |
| Linkage map scaffolded assembly | ERZ1089756 |
| PacBio reads | ERR3569680 - ERR3569745 |
| Dovetail reads | ERR3587405 - ERR3587406 |
| Male/Female reads | ERR3587407 -ERR3587409 |

## References

1. Kingan SB, Heaton H, Cudini J, Lambert CC, Baybayan P, Galvin BD, et al. A High-Quality De novo Genome Assembly from a Single Mosquito Using PacBio Sequencing. Genes. 2019;10:62.

2. Kajitani R, Yoshimura D, Okuno M, Minakuchi Y, Kagoshima H, Fujiyama A, et al. Platanus-allee is a de novo haplotype assembler enabling a comprehensive access to divergent heterozygous regions. Nat Commun. 2019;10:1702.

3. Roach MJ, Schmidt SA, Borneman AR. Purge Haplotigs: allelic contig reassignment for third-gen diploid genome assemblies. BMC Bioinformatics. 2018;19:460.

4. Pierini F, Lenz TL. Divergent Allele Advantage at Human MHC Genes: Signatures of Past and Ongoing Selection. Mol Biol Evol. 2018;35(9):2145–58.

5. Chin C-S, Peluso P, Sedlazeck FJ, Nattestad M, Concepcion GT, Clum A, et al. Phased diploid genome assembly with single-molecule real-time sequencing. Nat Methods. 2016 Oct;(October):1–7.

6. Kajitani R, Toshimoto K, Noguchi H. Efficient de novo assembly of highly heterozygous genomes from whole-genome shotgun short reads. Genome Res. 2014;24(8):1384–95.

7. Kronenberg ZN, Hall RJ, Hiendleder S, Smith TPL, Sullivan ST, Williams JL, et al. FALCON-Phase: Integrating PacBio and Hi-C data for phased diploid genomes. bioRxiv. 2018;1–12.

8. Koren S, Rhie A, Walenz BP, Dilthey AT, Bickhart DM, Kingan SB, et al. De novo assembly of haplotype-resolved genomes with trio binning. Nat Biotechnol. 2018;36:1174–82.

9. Malinsky M, Simpson JT, Durbin R. trio-sga: facilitating de novo assembly of highly heterozygous genomes with parent-child trios. bioRxiv. 2016;

10. Garg S, Aach J, Li H, Durbin R, Church G. A haplotype-aware de novo assembly of related individuals using pedigree graph. bioRxiv. 2019;

11. You M, Yue Z, He W, Yang X, Yang G, Xie M, et al. A heterozygous moth genome provides insights into herbivory and detoxification. Nat Genet. 2013 Mar;45(2):220–5.

12. Martins S, Naish N, Walker a S, Morrison NI, Scaife S, Fu G, et al. Germline transformation of the diamondback moth, Plutella xylostella L., using the piggyBac transposable element. Insect Mol Biol. 2012 Aug;21(4):414–21.

13. Li H. Aligning sequence reads, clone sequences and assembly contigs with BWA-MEM. 13033997v1 [q-bioGN]. 2013;

14. Walker BJ, Abeel T, Shea T, Priest M, Abouelliel A, Sakthikumar S, et al. Pilon: An Integrated Tool for Comprehensive Microbial Variant Detection and Genome Assembly Improvement. PLoS One. 2014;9(11):e112963.

15. Koren S, Walenz BP, Berlin K, Miller JR, Bergman NH, Phillippy AM. Canu: scalable and accurate long-read assembly via adaptive k -mer weighting and repeat separation. Genome Res. 2017;27:722–36.

16. Soderlund C, Nelson W, Shoemaker A, Paterson A. SyMAP: A system for discovering and viewing syntenic regions of FPC maps. Genome Res. 2006;16:1159–68.

17. Soderlund C, Bomhoff M, Nelson WM. SyMAP v3.4: a turnkey synteny system with application to plant genomes. Nucleic Acids Res. 2011;39(10):e68.

18. Thorvaldsdottir H, Robinson JT, Mesirov JP. Integrative Genomics Viewer (IGV): high-performance genomics data visualization and exploration. Briefings in BioinformaticsThoBa. 2012;14(2):178–92.

19. Laetsch DR, Blaxter ML. BlobTools: Interrogation of genome assemblies. F1000Research. 2017;6:1287.

20. Smit AFA, Hubley R, Green P. RepeatMasker Open-4.0 [Internet]. 2013. Available from: http://www.repeatmasker.org

21. Challis RJ, Kumar S, Dasmahapatra KK, Jiggins CD, Blaxter M. Lepbase: the Lepidopteran genome database. bioRxiv. 2016;

22. Huang S, Kang M, Xu A. HaploMerger2: rebuilding both haploid sub-assemblies from high-heterozygosity diploid genome assembly. Bioinformatics. 2017;33(16):2577–9.

23. Putnam NH, Connell BLO, Stites JC, Rice BJ, Blanchette M, Calef R, et al. Chromosome-scale shotgun assembly using an in vitro method for long-range linkage. Genome Res. 2016;26:342–50.

24. Baxter SW, Davey JW, Johnston JS, Shelton AM, Heckel DG, Jiggins CD, et al. Linkage mapping and comparative genomics using next-generation RAD sequencing of a non-model organism. PLoS One. 2011;6(4):e19315.

25. Catchen J, Hohenlohe PA, Bassham S, Amores A. Stacks: An analysis tool set for population genomics. Mol Ecol. 2013;22:3124–40.

26. Lunter G, Goodson M. Stampy: A statistical algorithm for sensitive and fast mapping of Illumina sequence reads. Genome Res. 2011;21:936–9.

27. Amores A, Catchen J, Nanda I, Warren W, Walter R, Schartl M, et al. A RAD-Tag Genetic Map for the Platyfish (Xiphophorus maculatus) Reveals Mechanisms of Karyotype Evolution Among Teleost Fish. Genetics. 2014;197:625–41.

28. Zimin A V, Puiu D, Luo M, Zhu T, Koren S, Marçais G, et al. Hybrid assembly of the large and highly repetitive genome of Aegilops tauschii, a progenitor of bread wheat, with the MaSuRCA mega-reads algorithm. Genome Res. 2017;27:1–6.

29. Kurtz S, Phillippy A, Delcher AL, Smoot M, Shumway M, Antonescu C, et al. Versatile and open software for comparing large genomes. Genome Biol. 2004;5(2):R12.

30. The International Silkworm Genome Consortium. The genome of a lepidopteran model insect, the silkworm Bombyx mori. Insect Biochem Mol Biol [Internet]. 2008 Dec;38(12):1036–45. Available from: http://www.ncbi.nlm.nih.gov/pubmed/19121390

31. Veeckman E, Ruttink T, Vandepoele K. Are we there yet? Reliably estimating the completeness of plant genome sequences. Plant Cell. 2016;28:1759–68.

32. Simão FA, Waterhouse RM, Ioannidis P, Kriventseva E V, Zdobnov EM. BUSCO: assessing genome assembly and annotation completeness with single-copy orthologs. Bioinformatics. 2015;31(19):3210–2.

33. Zdobnov EM, Tegenfeldt F, Kuznetsov D, Waterhouse RM, Ioannidis P, Seppey M, et al. OrthoDB v9.1: Cataloging evolutionary and functional annotations for animal, fungal, plant, archaeal, bacterial and viral orthologs. Nucleic Acids Res. 2017;45:D744--D749.

34. Derks MFL, Smit S, Salis L, Schijlen E, Bossers A, Mateman C, et al. The Genome of Winter Moth (Operophtera brumata) Provides a Genomic Perspective on Sexual Dimorphism and Phenology. Genome Biol Evol. 2015;7(8):2321–32.

35. Davey JW, Chouteau M, Barker SL. Major improvements to the Heliconius melpomene genome assembly used to confirm 10 chromosome fusion events in 6 million years of butterfly evolution. G3. 2016;6(March):695–708.

36. Zhan S, Merlin C, Boore JL, Reppert SM. The monarch butterfly genome yields insights into long-distance migration. Cell. 2011 Nov;147(5):1171–85.

37. Ahola V, Lehtonen R, Somervuo P, Salmela L, Koskinen P, Rastas P, et al. The Glanville fritillary genome retains an ancient karyotype and reveals selective chromosomal fusions in Lepidoptera. Nat Commun. 2014 Sep;5:4737.

38. Cong Q, Shen J, Borek D, Robbins RK, Otwinowski Z, Grishin N V. Complete genomes of Hairstreak butterflies, their speciation, and nucleo-mitochondrial incongruence. Sci Rep. 2016 Jan;6(April):24863.

39. Nowell RW, Elsworth B, Oostra V, Zwaan BJ, Wheat CW, Saastamoinen M, et al. A high-coverage draft genome of the mycalesine butterfly Bicyclus anynana. Giga Sci. 2017;6:1–7.

40. Kanost MR, Arrese EL, Cao X, Chen Y, Chellapilla S, Goldsmith MR, et al. Multifaceted biological insights from a draft genome sequence of the tobacco hornworm moth, Manduca sexta. Insect Biochem Mol Biol. 2016;76:118–47.

41. Mapleson D, Accinelli GG, Kettleborough G, Wright J, Clavijo BJ. Sequence analysis KAT: a K-mer analysis toolkit to quality control NGS datasets and genome assemblies. Bioinformatics. 2017;33(4):574–6.

42. Pinharanda A, Martin SH, Barker SL, Davey JW, Jiggins CD. The comparative landscape of duplications in Heliconius melpomene and Heliconius cydno. Heredity. 2017;118:78–87.

